# compaRe, an ultra-fast and robust suite for multiparametric screening, identifies phenotypic drug responses in acute myeloid leukemia

**DOI:** 10.1101/2021.01.08.425880

**Authors:** Morteza Chalabi Hajkarim, Ella Karjalainen, Mikhail Osipovitch, Konstantinos Dimopoulos, Sandra Leigh Gordon, Francesca Ambri, Kasper Dindler Rasmussen, Kirsten Grønbæk, Kristian Helin, Krister Wennerberg, Kyoung Jae Won

## Abstract

Multiparametric phenotypic screening of cells, for example assessing their responses to small molecules or knockdown/knockout of specific genes, is a powerful approach to understanding cellular systems and identifying potential new therapeutic strategies. However, automated tools for analyzing similarities and differences between a large number of tested conditions have not been readily available. Methods designed for clustering cells cannot identify differences between samples effectively. We introduce compaRe for ultra-fast and robust analysis of multiparametric high-throughput screening. Applying a mass-aware gridding algorithm using hypercubes, compaRe performs automatic and effective similarity comparison for hundreds to thousands of tests and provides information about the treatment effect. Particularly for screening data, compaRe is equipped with modules to remove various sources of bias.

Benchmarking tests show that compaRe can circumvent batch effects and perform a similarity analysis substantially faster than conventional analysis tools. Applying compaRe to high-throughput flow cytometry screening data, we were able to distinguish subtle phenotypic drug responses in a human sample and a genetically engineered mouse model with acute myeloid leukemia (AML). compaRe revealed groups of drugs with similar responses even though their mechanisms are distinct from each other. In another screening, compaRe effectively circumvented batch effects and grouped samples from AML and myelodysplastic syndrome (MDS) patients using clinical flow cytometry data.

## Introduction

Technological development has accelerated the generation of multiparametric cellular screening data through methodologies such as high-content microscopy and high-throughput flow cytometry in a large scale^1–3^. However, analyzing massive multiparametric datasets to provide an overview of similarity and differences between samples is still a challenge^1–3^. This analytical challenge is further complicated by various sources of bias often existing in the data such as batch effects and signal drift within a plate^1–3^.

There have been efforts to cluster samples from large-scale multidimensional screening data. A simple approach is to use a representative value for each cell marker such as median fluorescence intensity (MFI) for clustering samples^4^. However, a single representative value can easily lose the information about the biologically relevant variance within and between cell populations. Metaclustering approaches have been suggested to cluster samples based on the similarity of the centroids of subpopulations identified in the individual samples^5–7^. However, meta-clustering using centroids can be misleading when subclusters are not sufficiently distinct or the number of sub-clusters varies. The excessive computing cost also makes it hard for meta-clustering to be used for a large number of samples. Alternatively, predefined groups through manual gating or machine learning have been used to cluster samples^8, 9^. However, using prior knowledge is not suitable for *de novo* discoveries. Dimension reduction methods^10–12^ combined with the Jensen-Shannon divergence (JSD) metric have also been used to cluster multidimensional samples^11^. However, dimension reduction algorithms including factor analysis and principal component analysis still require excessive computing cost with an inherent information loss. It is also important to note that none of the methodologies developed so far efficiently correct for the sources of bias in multiparametric screenings.

To provide a useful tool for precise and effective interpretation of multiparametric screening data, we developed compaRe. compaRe measures the similarity between samples by dynamic gridding of the data. This mass-aware gridding increases the robustness of the algorithm to batch effect, signal drift and data size, while guaranteeing fast clustering performance as it does not require the computing costs for dimension reduction or subsampling. compaRe has also modules for quality control and bias correction which effectively reveal and remove various sources of bias in the screening data. compaRe provides various visualization tools that help with the interpretation of the results.

We show that compaRe can effectively circumvent batch effects in multidimensional screening data while showing remarkably fast clustering. Using compaRe to analyze high-throughput flow cytometry screening of AML cells, we successfully removed various sources bias and grouped drugs based on their responses. compaRe effectively revealed that groups of drugs showed similar responses even though their known mechanisms are different from each other. compaRe was sensitive enough to capture subtle changes in drug responses. We further applied compaRe to 25 clinical flow cytometry data sets representing AML and MDS patients. Without prior knowledge, compaRe could effectively group the samples based on the disease with indications that the grouping also linked to clinical outcome. Thus, we conclude that compaRe is a useful tool for exploring complex multiparametric datasets and to help interpretation of drug responses.

## Results

### compaRe is a tool for clustering and visualizing multiparametric screening data

compaRe is designed to analyze multiparametric screening data from small screenings of just a few samples to high-throughput screenings of hundreds of samples. At the core of compaRe is a module for measuring similarity between samples using dynamic gridding (**Figure 1, Online Methods**). Given two samples, the dynamic gridding algorithm divides a high-dimensional space (formed by for example fluorescence markers) of each sample into a number of spatial units called hypercubes. The proportion of data points for the corresponding hypercubes of each sample is used to obtain the similarity. In this setting, compaRe becomes robust to the difference in the data size, presence of batch effect and signal drift.

**Figure 1:**
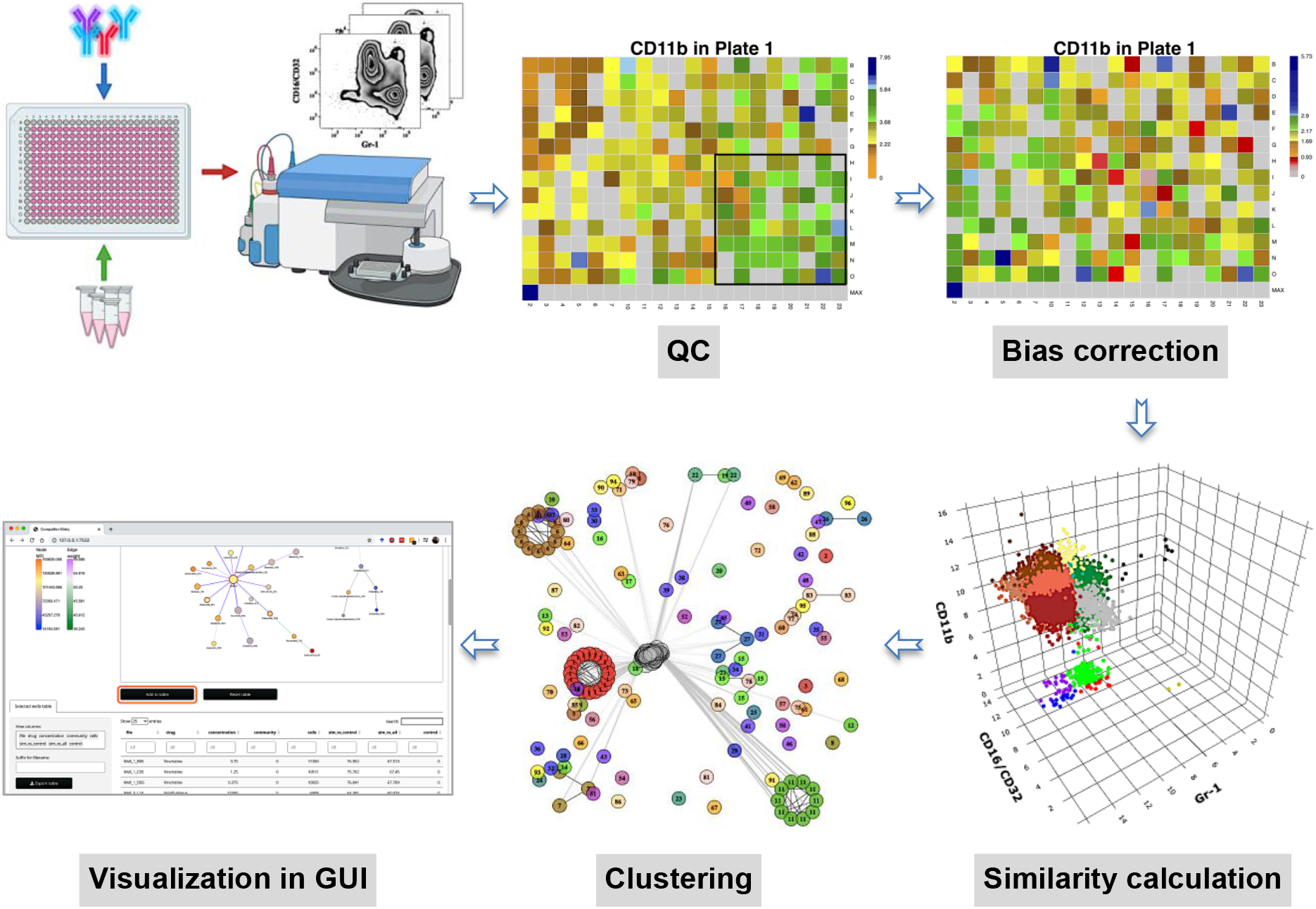
compaRe is a comprehensive suite for multiparametric screening data. High-throughput flow cytometry generates massive multidimensional data from hundreds of samples. compaRe’s **quality control (QC)** module reveals several sources of bias in the assay such as signal (intensity difference between the top left and bottom right corners) and cell viability drifts. These two are corrected for in the **bias correction** modules within and between the plates. compaRe performs a pairwise **similarity calculation** between the samples using dynamic gridding and forming hypercubes (represented by distinct colors). The portions of the data within individual hypercubes are used to calculate similarity. **Clustering** is performed based on the similarity. The **graphical user interface (GUI)** provides various ways to thoroughly explore and visualize the read-outs.

Besides its core module, compaRe comprises several modules for quality control, bias correction, clustering and visualization. **Figure 1** shows the modules for a high-throughput flow cytometry drug screening of mouse AML cells. During quality control, several sources of bias in screening data such as autofluorescence, bioluminescence, carryover effect, edge effect, signal drift and the drift in the number of live cells (cell viability drift) are identified. Samples and even an entire microtiter plate which are worst affected by like autofluorescence and carryover effect can be discarded at this point. The bias correction modules effectively correct for signal and cell viability drifts, the two main sources of bias, using regression analysis (**Figure 1, Online Methods**). Based on similarity, compaRe performs clustering using a graphical algorithm (**Figure 1, Online Methods**). Initially, all nodes (samples) are connected forming a complete weighted graph wherein weights represent similarity values. The graph is then pruned to remove potential false positive edges using a threshold inferred from negative controls. After constructing a linked graph, clustering is tantamount to finding maximal cliques (complete subgraphs that cannot be extended) each containing similar samples like drugs with similar responses.

compaRe carries out the analysis using parallel computing. Modules can also run independently of each other. The similarity and clustering modules of compaRe can be applied to any other tabular datasets.

### COMPARE’s similarity comparison is ultra-fast and robust to batch effect

To evaluate the performance of compaRe’s similarity module, we tested compaRe, JSD with UMAP and meta-clustering with PhenoGraph^6^ on a set of mass cytometry data containing phenotypically heterogeneous (AML patients) and homogeneous (healthy donors) bone marrow aspirates^6^. The set included 21 samples labeled with 16 surface markers collected from 16 pediatric AML patients and 5 healthy adult donors. We introduced random noise with Gaussian distribution to the 16 parameters of each sample to simulate batch effect. In this setting, although the added noise undermines the similarities, the overall cell population morphology is not altered, and the simulated samples will still have the highest similarity with their original counterparts.

Even with the added noise, compaRe precisely retrieved the original counterparts as the most similar ones (**Figure 2a**). However, the batch effect seriously compromised the performance of metaclustering and JSD, showing a number of maximum similarities other than the originals (**Figure 2b-c**). These results demonstrate the advantages of using dynamic gridding for the comparison between samples with batch effect.

**Figure 2:**
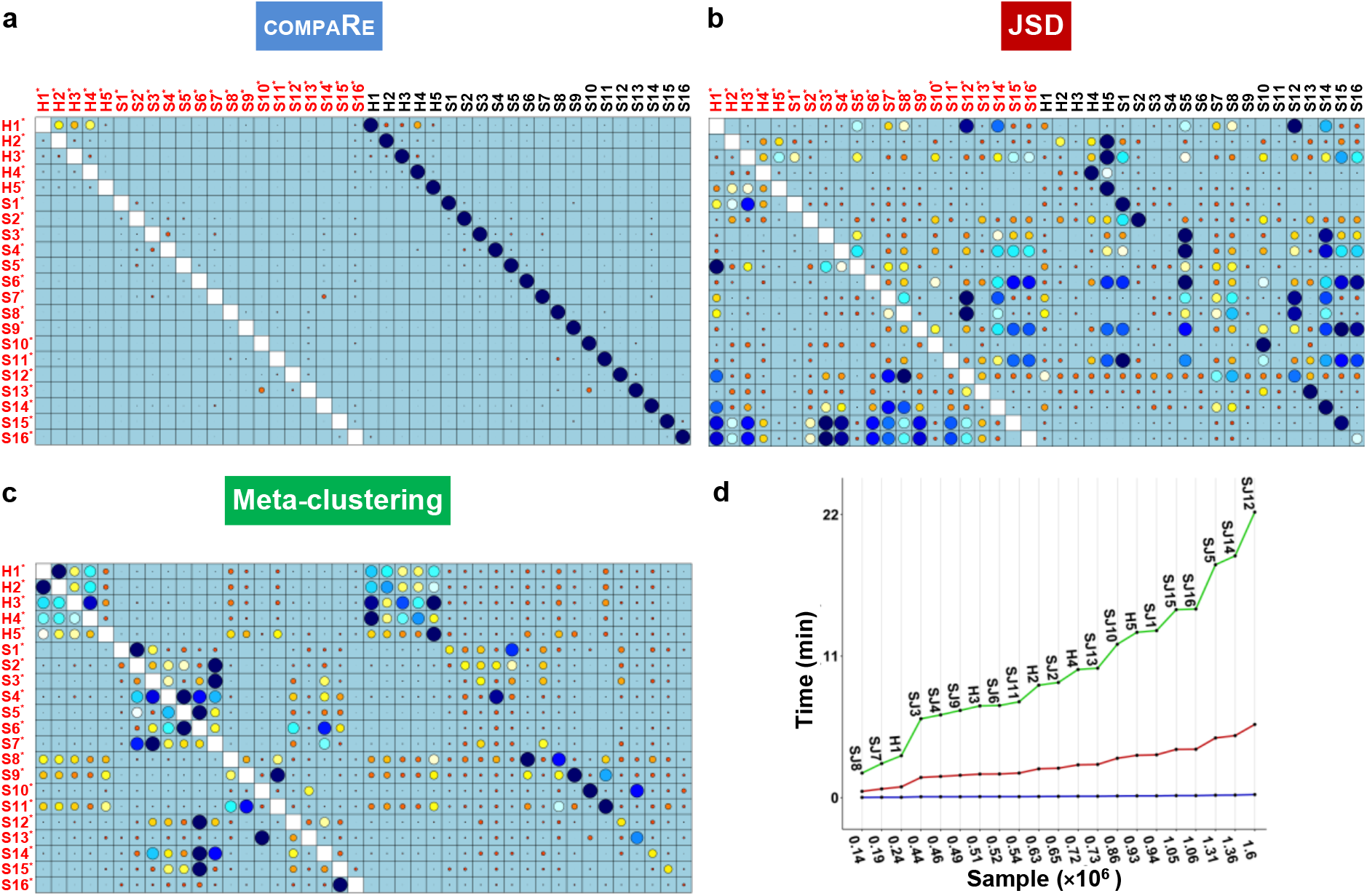
compaRe robustly measures the similarity between samples in the presence of batch effect. Similarity matrix generated by compaRe is shown in **(a)**. Size and color of dots represent the level of similarity. Self-comparisons were removed. Noise was added (marked with *) to the original 21 mass cytometry samples of bone marrow aspirates from 16 pediatric AML patients (S) and 5 healthy adult donors (H). Similarity matrices using JSD with UMAP and meta-clustering with PhenoGraph are shown in **(b)** and **(c)** respectively. **(d)** shows the run time of comparing each sample to itself. Samples were sorted based on their size.

Additionally, we investigated the run time of the three methods (on a computer server with 2.30 GHz CPU and 256 GB DDR4 memory) by comparing each sample to itself (**Figure 2d**). The running time increased steeply for meta-clustering and JSD as the sample size increased. Compared with metaclustering and JSD, the increase of computing time for compaRe is almost unnoticeable. Applied to all of the 21 samples, compaRe took only 25 min to finish the analysis for the 210 pairwise comparisons (this is without subsampling or dimension reduction of the data and the risk of losing rare subpopulations), while meta-clustering and JSD took 39 h and 10 h respectively. For the feasibility of JSD with UMAP, we subsampled each sample to 100,000 cells (default value suggested in^11^). When we fixed this limit to 60% of each sample, the computing time of JSD increased to 3 days.

### Identifying cell subtype-specific drug responses in high-throughput flow cytometry of mouse AML cells

We applied compaRe to high-throughput flow cytometry data to identify cell subtype-specific responses evoked by antineoplastic agents in leukemic spleen cells from an AML mouse model. Splenic cells were sorted for c-Kit cell surface expression, allowing for the enrichment of stem/progenitor-type leukemic cells. On *ex vivo* expansion, these cells differentiate and expand in a similar way as *in vivo* with a clear stem cell/progenitor population and partial differentiation towards CD11b/Gr-1 or CD16/CD32-expressing differentiated myeloid cells. After *ex vivo* expansion, the leukemic cells were plated onto multi-well plates containing a library of 116 antineoplastic agents including surface and nuclear receptor inhibitors and activators, enzyme inhibitors and cytotoxic chemotherapy in a five-point concentration range as well as 20 negative control wells (**Supplementary Table 1**). After 72 h of drug exposure, we stained the cells with fluorescently labeled antibodies against three cell surface markers (CD16/32, Gr-1 and CD11b) and quantified cell surface marker expression using a high-throughput flow cytometer.

compaRe corrected the signal drift and sources of bias in cell numbers, as well as inter-plate sources of bias (**Supplementary Figure 1**). After clustering and clique analysis, we obtained 134 cliques, each sharing similar drug responses (**Supplementary Table 2**).

To get an overview of the assay, we generated a dispersion map of the clusters (**Figure 3a-b** and **Online Methods**). We identified a distinct response group characterized with decreased Gr-1 and concomitant increase of CD16/CD32 as compared to control (Group 1 in **Figure 3a**). Most of the cliques included in this response group consisted of drugs in high concentrations with cytotoxic/cytostatic effects. However, some drugs in this group had a milder effect on live cell numbers, and these were enriched for mitogen-activated protein kinase (MAPK) pathway-associated inhibitors (**Figure 3c, Supplementary Table 3**). For instance, trametinib (2.5 nM) in clique 23 (C23) showed marked decrease of Gr-1 and increase of CD16/CD32, further confirming the results of compaRe (**Figure 3d**). The MAPK pathway is a regulator of diverse cellular processes such as proliferation, survival, differentiation and motility^13^. Our findings suggest that MAPK signaling controls the differentiation and/or proliferation towards Gr-1-/CD16+ cells.

**Figure 3:**
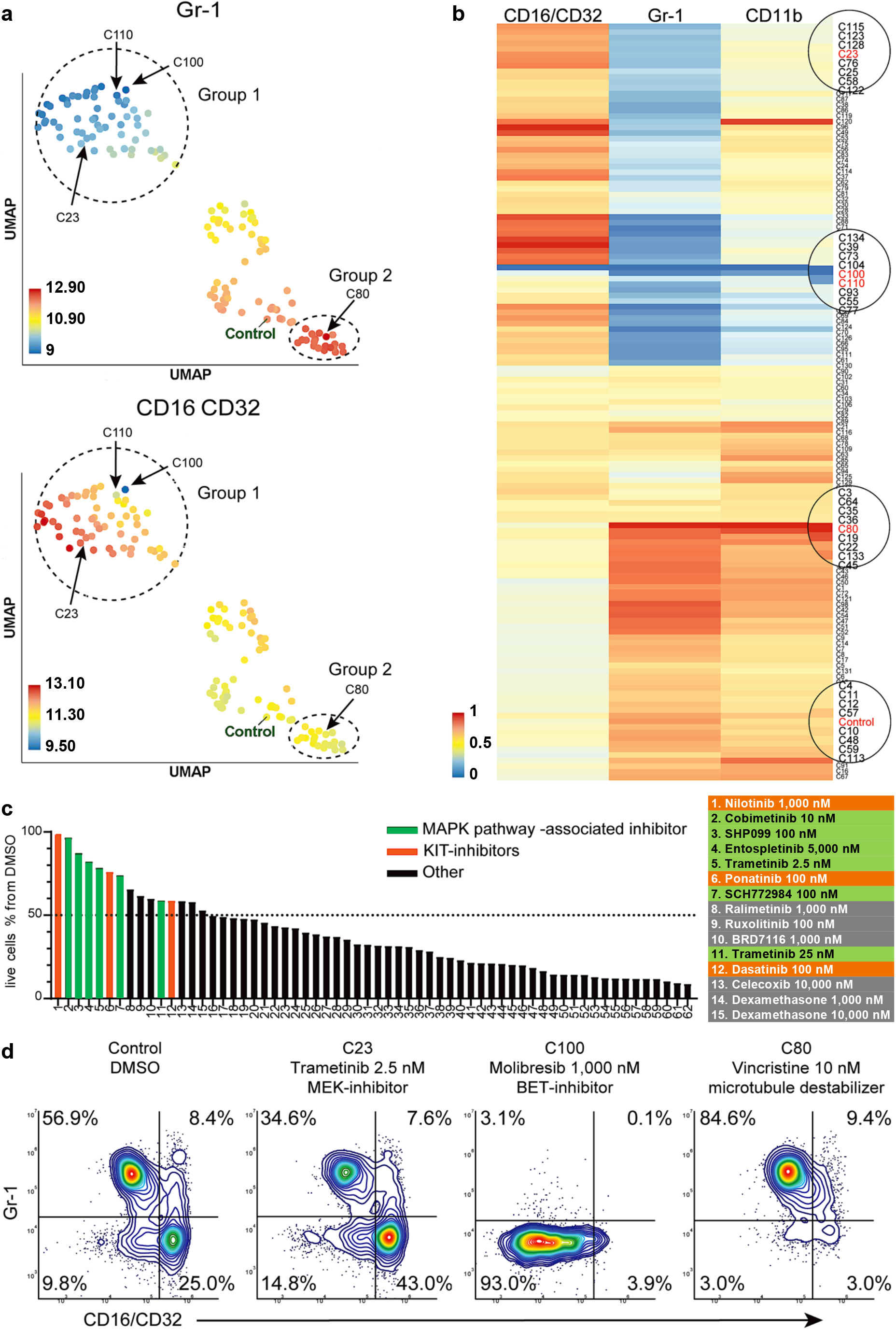
compaRe analysis identifies several distinct cell subtype-specific responses in a high-throughput flow cytometry screening of mouse AML cells. **(a)** A UMAP plot of cliques identified by compaRe. Cliques are colored by Gr-1 and CD16/CD32 MFIs. Group 1 is characterized with reduced Gr-1 and increased CD16/CD32 as compared with control. Group 2 has increased Gr-1 expression compared with control. (**b**) Heatmap of marker MFIs. Values are normalized between 0 and 1 per marker to make cross-comparisons possible. Cliques containing control, trametinib (2.5 nM) (C23), molibresib and birabresib (C100 and C110), and vincristine (C80) are marked. (**c**) Waterfall plot of compounds belonging to response group 1, showing live cell count as a percentage of control treatment (DMSO). (**d**) Density scatter plots for Control (DMSO), C23, C100, and C80.

In high concentration, molibresib and birabresib, inhibitors of BET proteins BRD2, BRD3, and BRD4, caused a reduction in live cell counts but also a reduction of MFI in all the measured markers, which corresponds to the loss of differentiation marker positive cells (Gr-1+, CD11b+, CD16/CD32 high) (**Figure 3b**: C100, C110, **Figure 3d**). The BRD2/3/4 proteins regulate transcription via recognition of acetylated lysines on histones and concomitant recruitment of other transcription and chromatin remodeling factors to enhance transcriptional activity^14^. The enrichment of seemingly highly undifferentiated cells could therefore be due to an early block in differentiation or that inhibition of BRD2/3/4 has led to a general decrease of cell surface protein transcription.

In this cell model, the leukemic stem-like cells are expected to be present within the differentiation marker negative population. These cells are potential targets for the treatment against leukemia. We observed response group 2 (**Figure 3a**) had a higher MFI in marker Gr-1 as compared to control, which could indicate a higher proportion of marker positive cells and thus, a lower proportion of marker negative ones. For most treatments in group 2, the increase in Gr-1 MFI was very slight and seemed to be linked to toxic drug concentrations. However, three drugs, vincristine (C80), tazemetostat, and tretinoin clearly reduced the proportion of differentiation marker negative cells (**Figure 3d**). Interestingly, these three drugs have very different modes of action: vincristine is a microtubule polymerization inhibitor, tazemetostat inhibits the histone methyltransferase EZH2, and tretinoin is a retinoic acid receptor agonist (**Supplementary Table 3**).

Taken together, compaRe analysis of the high-throughput flow cytometry screening allowed rapid identification of several distinct phenotypic responses in this mouse AML model, as well as the cellular signals that drive them.

### Identifying a set of drugs that induce expansion of a distinct CD34+/CD38+ cell population in an AML patient

We further applied compaRe to the drug screening data from AML patient samples. Primary AML bone marrow mononuclear cells were dispensed into a 384-multiwell plate containing a library of 40 drugs and drug combinations in 7-point concentration ranges (**Supplementary Table 4**). After 72 h of drug exposure, the cells were stained with fluorescently labeled antibodies against a panel of AML-related cell surface markers (CD45, CD34, CD38, CD117, HLA-DR, CD45-RA, CD3 and a mix of myeloid differentiation-related markers). A high-throughput flow cytometer was used to quantify cell surface marker expression.

compaRe analysis identified several distinct response groups (**Figure 4a, Supplementary Table 5**). Response group 1 had notably higher MFIs in the CD34 and CD38 channels compared to controls. Interestingly, the increase in MFIs was due to a drug concentration-dependent appearance of a CD34+/CD38+ cell population that was barely detectable in the DMSO control samples (**Figure 4b**). The appearance of this CD34+/CD38+ population was also concomitant to a general increase in live cell count (**Figure 4c**). Altogether, seven different drugs had the same effect (**Figure 4d**), most of them being highly selective signal transduction inhibitors such as trametinib (MEK inhibitor), copanlisib (PI3K inhibitor) and PIM447 (PIM kinase inhibitor).

**Figure 4:**
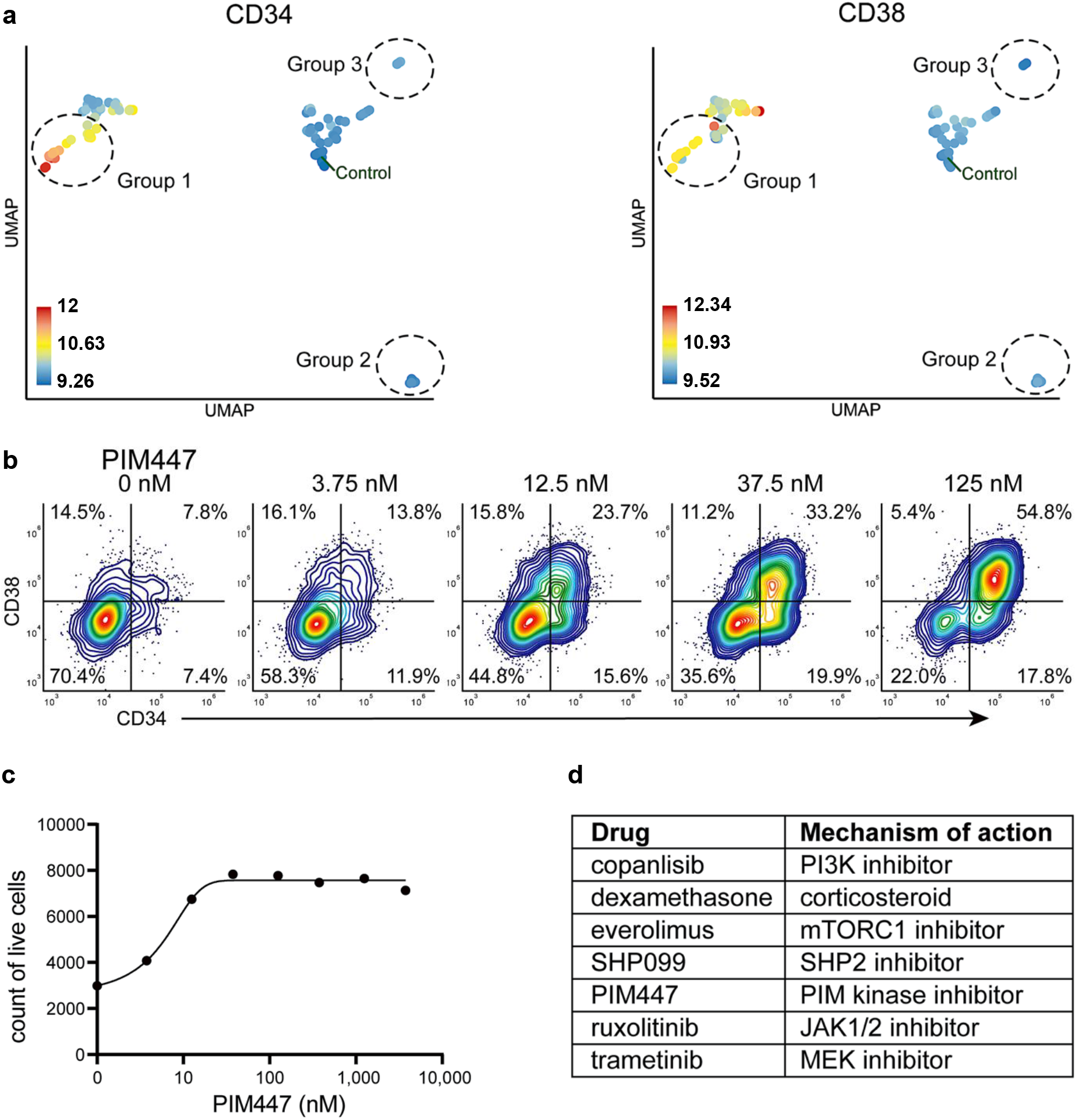
Identification of drugs which induce expansion of CD34+/CD38+ cells in an AML patient. (**a**) UMAP of cliques identified by compaRe. Cliques are colored by CD34 and CD38 MFI. Response groups of interest are indicated using a dashed line. (**b**) Example of response group 1: density scatter plot of markers CD34 and CD38 in different concentrations of PIM kinase inhibitor PIM447. (**c**) Count of live cells after 72 h exposure to different concentrations of PIM kinase inhibitor PIM447. (**d**) Table of drugs which induced expansion of the CD34+/CD38+ cell population.

Response group 2 consisted of two drugs: birabresib and lenalidomide in different concentrations. These induced a decrease in MFI in the CD45-RA and CD45 channels (**Supplementary Figure 2a**). In the case of lenalidomide this response was likely due to cell toxicity and/or growth inhibition (**Supplementary Figure 2b**). Interestingly, birabresib response was very pronounced without the loss of live cell numbers, (**Supplementary Figure 2b**) but with a decrease in MFI in the channel for the cell differentiation marker mix (**Supplementary Figure 2c**). compaRe detected response group 3 was distinct from the controls. This group includes treatment with tretinoin (several concentrations), navitoclax, and mitoxantrone (low dose).

Further validation showed the phenotypic response in group 3 is subtle but with a distinct increase in CD34+ cells (**Supplementary Figure 2d**). This result highlights that compaRe analysis is sensitive enough to identify these subtle changes.

### Immunophenotypic similarities measured by compaRe correlate with MDS and AML patients’ prognosis

MDS is caused by dysplastic hematopoietic stem cell-like cells which suppress normal hematopoiesis and initially results in cytopenia, which later on presents with more severe symptoms due to clonal progression and the lack of a functional hematopoiesis. In addition, approximately one third of MDS cases progress to AML^15, 16^. MDS is typically assigned a risk level based on the number of immature blood cells in the bone marrow, level of cytopenia, and the presence of known poor prognosis cytogenetic alterations. MDS is incurable except for allogeneic bone marrow transplantation, which is not an option in the majority of patients due to high age and comorbidities. Cases scored as low risk (LR-MDS) tend to progress slowly. Most cases of LR-MDS in the elderly are managed with growth factors and transfusions to maintain red blood cell counts. In contrast, high-risk MDS (HR-MDS) is likely to progress quickly or transform to AML and survival is expected to be short and more intensive treatment, including hypomethylating agents, is often applied. Therefore, it is important to weigh the benefits of treatment options against the possible risks and side effects.

We analyzed the clinical flow cytometry data from bone marrow mononuclear cell samples taken at diagnosis for 12 LR-MDS patients, 4 HR-MDS patients (as determined by the revised international prognostic scoring system (IPSS) total risk score^17^) and 9 AML patients, collected between November 2015 and November 2018. The sample cohort was blindly analyzed with compaRe using the data from a standardized clinical flow cytometry panel detecting 20 markers in 4 different separately stained tubes (**Supplementary Table 6**). Often, flow cytometry data are difficult to standardize between batches analyzed days or months apart, as the cytometer’s settings can change with time, or reagents may fade. The batch effect on this data set was inevitable due to a 3-year-time span of collection and analysis. However, since compaRe is able to circumvent batch effect, we were able to apply it to assess similarity between the samples.

We could identify immunophenotypically similar patients and, as expected, MDS samples were overall more similar to each other than the AML samples, which showed a greater heterogeneity (**Figure 5a**). In comparing which samples showed greatest similarity to a given sample together with information about clinical outcome (**Figure 5b, Supplementary Table 7**), it appeared that samples from cases with a good clinical outcome most closely resembled other good outcome cases and pooroutcome cases resembled other aggressive cases. For example, HR-MDS-1 and −2, where the diagnostic samples had very similar immunophenotypic profiles, both progressed to AML. Also, patient HR-MDS-4 whose sample showed a high similarity to an AML patient’s sample progressed to AML. Conversely, patient AML-2, whose sample resembled an LrMDS profile upon data collection, went into remission with only one cycle of induction chemotherapy. For the AML samples, even though they were heterogeneous, the highest similarities were seen with cases with similar patient outcomes. Overall, these results show that compaRe can be used to analyze similarities between complex sets of clinical flow cytometry data with batch effects and that the analysis could potentially be informative for prognostic purposes, which however will require analyses of larger cohorts of uniformly treated patients.

**Figure 5:**
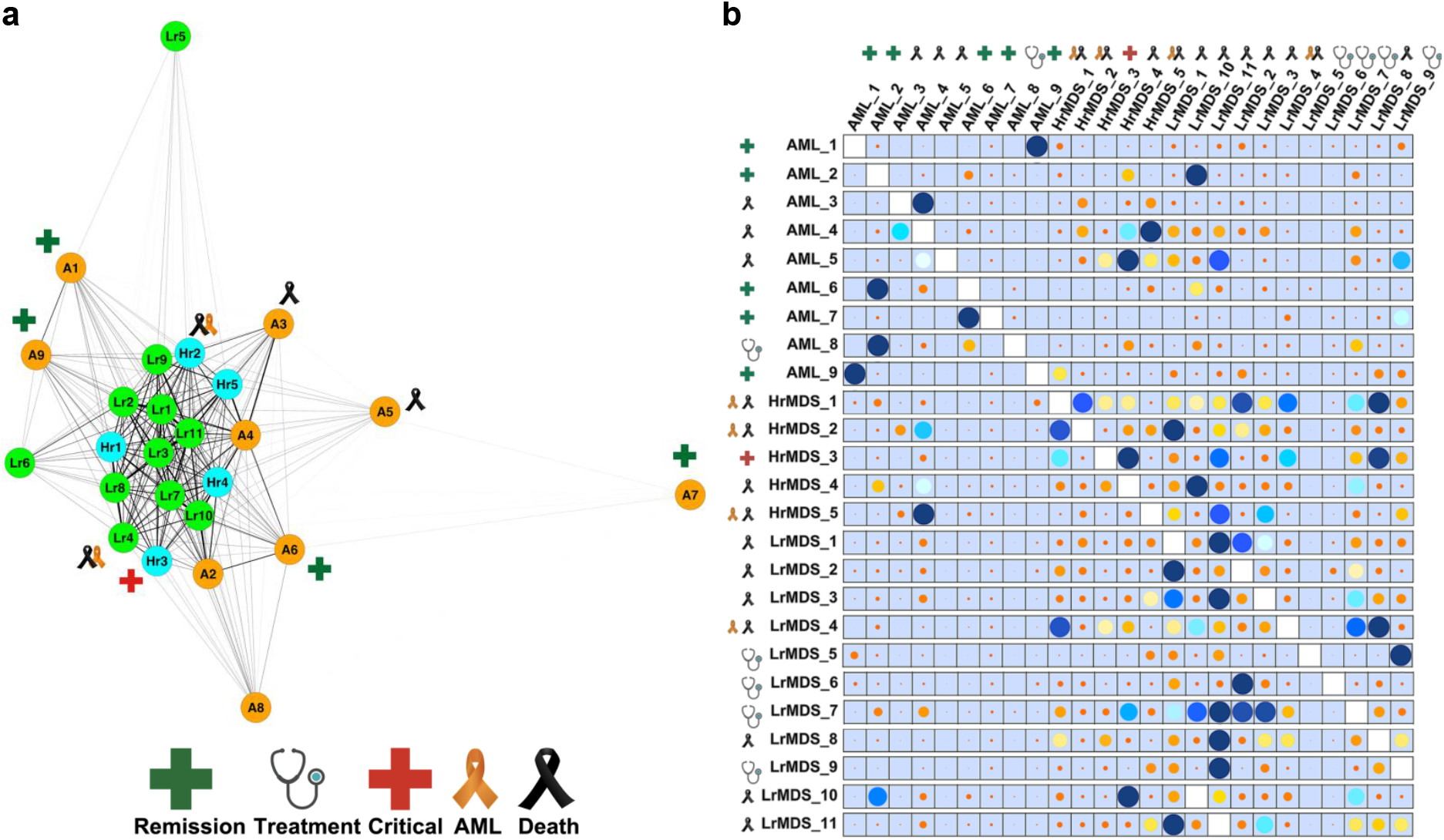
Immunophenotypic similarities measured by compaRe correlate with prognosis of AML and MDS patients. (**a**) The similarity graph. MDS samples formed a dense community in the middle with AML patients around them. LR-MDS-5 and AML-7 showed low similarity with the rest of the samples. **(b)** The similarity table. For each row, similar samples were marked with the enlarged size and darker color of the dots. Values in each row were normalized to accentuate the dynamic range. The table legend shows the clinical conditions of patients at the time of writing.

## Discussion

Technological developments in phenotypic screening have enabled testing thousands of conditions in parallel. Algorithmic development is essential to group the samples based on the cell population change. We developed compaRe to analyze this type of multidimensional phenotypic screening data. compaRe is unique in many ways compared to previous analysis methods. compaRe utilizes a dynamic gridding algorithm, which substantially reduces the computing costs and provides robustness to batch effects and signal drift in the input data. Compared with it, meta-clustering and JSD requires a considerable amount of time for clustering or obtaining probability distribution.

We tested the effectiveness of dynamic gridding by checking the similarity after adding noise. compaRe effectively circumvented the batch effect and found the best match with the original counterparts while JSD and meta-clustering (with PhenoGraph) analysis suffered from the batch effect. The poor accuracy of meta-clustering demonstrates the drawbacks of using the centroids for similarity comparison across samples. It is of note that compaRe did not have to subsample the data and ran more than 20 times faster than JSD for analyzing mass cytometry data from 21 patients. Likewise, strategies to cluster a handful of multi-dimensional data such as CyTOFmerge^18^ cannot be applied for large-scale screening data. Our results indicate that approaches such as meta-clustering cannot be applied to multiparametric screening that generates data from hundreds to thousands of conditions. It was indeed impossible for us to benchmark compaRe to other meta-clustering approaches with screening data due to their exhaustive computing cost.

compaRe is equipped with several modules for quality control, bias correction and visualization. compaRe has an option to correct for signal and cell viability drifts when needed making it suitable to analyze long run screening data. In our explored datasets, signal drift was obviously associated with the order in which wells were read. It was caused by the time differences in antibody incubation across the plate as the flow cytometer requires more than one hour to sample all wells in a 384-well plate. For high-density assay plate formats with a large number of wells, this can cause gradual incremental influences in intensity and cell viability. Therefore, when aligning wells along the order that the flow cytometer sampled the wells, we found a linear trend in MFIs. We benefited from regression analysis to remove the effect of signal drift.

compaRe provides guidance in interpreting drug responses with its visualization tools. We identified similar responses evoked by different doses of antineoplastic agents in high-throughput flow cytometry screening of an AML mouse model and an AML human patient. In the mouse screening, the results allowed us to identify key signals in the cells that control different steps of differentiation. We also identified drugs with different mechanisms exhibiting similar responses, which may suggest crosstalk between different signaling pathways. Importantly, we could also identify subtle but distinct phenotypic drug-induced changes. During the analysis, compaRe made it easy to explore a very complex dataset and was able to identify multiple cell-subtype-specific drug responses in a substantially reduced timeline.

Using compaRe, we evaluated the immunophenotypic similarity of 25 AML and MDS patient samples in a clinical flow cytometry data set collected upon diagnosis. It is encouraging that this pilot analysis indicates compaRe can generate clinically relevant information. The assessment of compaRe similarity analysis in relation to clinical outcome showed that AML patients whose immunophenotypic profiles resembled profiles of non-aggressive MDS cases were associated with a good outcome. Conversely, MDS samples, showing similarity with aggressive AML profiles, were associated with a poor outcome, including transformation into AML. We speculate that analysis of clinical flow data by compaRe could potentially be a prognostic tool in hematological malignancies, either as an alternative or a supplement to conventional risk assessments.

compaRe can entirely run through the command-line interface on computer servers or the graphical user interface (GUI) on desktops. The GUI provides the investigator with numerous interactive visualization tools including cell staining, graphical representation and gating. In sum, compaRe provides a total package for fast, accurate and readily interpretable multiparametric screening data analysis.

## Online Methods

### Mass cytometry of healthy and pediatric AML bone marrow aspirates

Mass cytometry dataset for 21 samples labeled with 16 surface markers collected from 16 pediatric AML patients obtained at diagnosis and 5 healthy adult donors^6^ were downloaded from Cytobank with the experiment ID <u>44185</u>. There are 378 FCS files in this experiment with one FCS file for each of 21 patients for each of 17 conditions (2 basal replicates and 16 perturbations). All FCS files from a single patient had been pooled then clustered with the PhenoGraph algorithm. Each file includes a column named PhenoGraph that specifies the PhenoGraph cluster to which each event was assigned as an integer. A value of 0 indicates no cluster was assigned because the cells were identified as outliers during some stage of analysis. Using the PhenoGraph column, we determined centroids of cell clusters, and used PhenoGraph to metacluster them as described in Lavine *et al*^6^. To generate the similarity matrix, we adapted an approach similar to that of compaRe so that each meta-cluster as a spatial unit was treated like a hypercube. We set compaRe’s *n* to 4 for this data set (**Online Methods** and **Supplementary Notes**).

### High-throughput flow cytometry of AML mouse model

AML primary splenic cells from *Npm1*^+/cA^; *Flt3*^+/ITD^; *Dnmt3a*^+/-^; Mx1-Cre+ moribund mice were sorted for c-Kit positivity and expanded *ex vivo*. AML cells were treated with a library of 116 chemotherapy and immunotherapy antineoplastic agents in a five-point concentration range (**Supplementary Table 1**). Treated samples were stained with three informative cell surface antibodies **(Supplementary Table 8)** and fluorescence was detected using a high-throughput flow cytometer iQue Screener Plus (Intellicyt). We set compaRe’s *n* to 5 for this assay.

### High-throughput flow cytometry of an AML human patient sample

Mononuclear cells were isolated from a donated human bone marrow aspirate from an AML patient (Danish National Ethical committee/National Videnskabsetisk Komité permit 1705391). The cells were treated with a library of 40 chemotherapy and immunotherapy antineoplastic agents in a seven-point concentration range (**Supplementary Table 4**) for 72 h. Cells were subsequently incubated with fluorescently labeled antibodies targeting 11 informative cell surface proteins in 8 fluorescence channels (**Supplementary Table 9**). Samples were read using a high-throughput flow cytometer (iQue Screener Plus, Intellicyt). We set compaRe’s *n* to 3 for this assay.

### Flow cytometry of AML and MDS patients

Clinical flow cytometry data using a slightly modified AML panel as described by the Euroflow Consortium^19^ from 25 bone marrow aspirates from MDS and AML patients from Rigshospitalet (Copenhagen, DK) were used for analysis. Each sample was analyzed using a total of four tubes (Euroflow AML panel tubes 1-4) with eight antibodies in each tube (**Supplementary Table 6**). Acquisition of data was performed on a FACS Canto (Becton Dickinson Immunocytometry Systems) and data analysis was done in the Infinicyt software (Cytognos, Salamanca, Spain). We set compaRe’s *n* to 5 for this assay.

### Quality control (QC)

Microplate heatmaps of medians come in handy in QC to reveal possible issues such as signal and cell viability drifts occurring during screening. However, as a typical heatmap has an equally spaced color palette, small but significant differences between wells are obscured and probably not visible. Therefore, we normalized the color palette by the distribution of the medians. Also, before clustering, we removed outliers in the negative controls that were different from the others in terms of similarity values measured by compaRe.

### Correcting signal and cell viability drifts

Depending on the protocol by which wells are processed, time may become a major concern so that some specific wells may have lower or higher values than expected. To correct for these sources of bias, we employed a two-step correction: intra-plate drift correction and inter-plate drift (batch effect) correction. For a given plate, we first fit a linear regression model and then vertically translate points (well values) with respect to the learned line as it rotates to the slope zero. After correcting for the intra-plate bias, the inter-plate bias is corrected by aligning medians of the plates, that is, translating to a common baseline.

### Similarity calculation using dynamic gridding

To measure the similarity between two datasets, compaRe divides each dimension into *n* subsets for each dataset individually so that a dataset with *d* dimensions (markers) will be gridded into at most *n^d^* spatial units called hypercubes. compaRe grids only the part of the space encompassing data points, avoiding empty regions. It then measures the proportion of data points for either dataset within each of the corresponding hypercubes. The difference between the two proportions is indicative of the similarity within that relative spatial position represented by each hypercube. The similarity in the exclusive hypercubes is considered 0. Averaging these differences across all the hypercubes indicates the amount of similarity between the two datasets.

compaRe captures the morphology of data enabling it to measure similarity even without correcting for signal drift or batch effect (**Supplementary Notes**). This way, two technical replicates analyzed by two different instruments or configurations suffering from signal shift will still have the highest similarity. To generate a similarity matrix of multiple input samples, compaRe runs in parallel. The similarity matrix could then be used for identifying clusters of samples such as drugs with similar dose responses.

### Graphical clustering of samples

To cluster samples, we developed a graphical clustering algorithm in which initially all nodes (samples) are connected forming a weighted complete graph wherein edges represent similarity between nodes. This graph is then pruned to remove potential false positive edges for a given cutoff inferred from negative controls. The optimal cutoff turns out to be the minimum weight in the maximum spanning tree of negative control nodes. After pruning, some samples may end up being connected to the negative controls (biologically inactive agents) and some disconnected (active agents). After constructing this graph, clustering is tantamount to finding maximal cliques among potent agents. In addition to maximal cliques, it also reports communities (a clique is actually a subset of a community). Communities can be seen as loose clusters. In a community, unlike a clique, similarity is not necessarily transitive meaning that if A is similar to B and B is similar to C, A is not necessarily similar to C. If these were three drugs within a community, concluding they had an equal response was not necessarily right unless they would form a clique.

### Dispersion graph and Dispersion map

compaRe visualizes the similarity of samples in the form of a dispersion graph by constructing their maximum spanning tree (**Supplementary Notes**). compaRe also uses UMAP to represent a dispersion map of clusters. The map is constructed using the centroid (median) of each clique. An informative map shows different groups by coloring the centroids according to their value. These groups are mostly the identified communities the cliques come from.

## Data Availability

Human AML high-throughput flow cytometry data have been deposited in FLOWRepository with the repository ID FR-FCM-Z3DP. Mouse AML high-throughput flow cytometry data have been deposited in FLOWRepository with the repository ID FR-FCM-Z357.

## Code Availability

Acquisition, installation and more technical details are available in the compaRe’s online tutorial on GitHub. Similarity measurement and clustering modules as stand-alone tools have been merged into a separate R package and are available for download at GitHub.

## Acknowledgement

This work was supported through a center grant from the Novo Nordisk Foundation (Novo Nordisk Foundation Center for Stem Cell Biology, DanStem; Grant Number NNF17CC0027852) and is also part of the Danish Research Center for Precision Medicine in Blood Cancers funded by the Danish Cancer Society (Grant number R223-A13071) and Greater Copenhagen Health Science Partners.

## Author Contributions

KW, KJW, MCH and EK conceived, designed and wrote the study. MCH developed the suite and analyzed the data. EK performed and analyzed flow cytometry screening experiments and evaluated the suite and the GUI. MO developed the GUI. SLG and FA designed and completed the AML patient sample drug screening. KDR and KH generated the mouse cell models. KK and KG assembled and annotated clinical flow cytometry data and assisted in its analysis. The manuscript was approved by all authors.

